# A single droplet digital PCR for *ESR1* activating mutations detection in plasma

**DOI:** 10.1101/507608

**Authors:** Emmanuelle Jeannot, Lauren Darrigues, Marc Michel, Marc-Henri Stern, Jean-Yves Pierga, Aurore Rampanou, Samia Melaabi, Camille Benoist, Ivan Bièche, Anne Vincent-Salomon, Radouane El Ayachy, Aurélien Noret, Nicolas Epaillard, Luc Cabel, François-Clément Bidard, Charlotte Proudhon

**Affiliations:** Circulating tumor biomarkers laboratory, Inserm CIC 1428, Institut Curie, PSL Research University, Paris, France; Department of Biopathology and Genetics, Institut Curie, PSL Research University, Paris, France; INSERM U830, Institut Curie, PSL Research University, Paris, France; Department of Medical Oncology, Institut Curie, PSL Research University, Paris & Saint Cloud, France; Paris Descartes University, Paris, France; Versailles Saint Quentin University, Paris Saclay University, Saint Cloud, France

**Author notes:** These authors share senior co-authorship (François-Clément Bidard, Charlotte Proudhon). **Corresponding authors:** Charlotte Proudhon, Institut Curie, 26 rue d’Ulm, 75005 Paris, France.

**Keywords:** *ESR1* mutation, ddPCR, circulating tumor DNA, breast cancer, monitoring

## Abstract

**Background:** Activating mutations in the estrogen receptor 1 (*ESR1*) gene are recurrent mechanisms of acquired resistance to aromatase inhibitors (AI), and may be the target of other selective estrogen receptor down-regulators. To assess the clinical utility of monitoring *ESR1* resistant mutations, a droplet digital PCR (ddPCR)-based assay compatible with body fluids is ideal due to its cost-effectiveness and quick turnaround.

**Methods:** We designed a multiplex ddPCR, which combines a drop-off assay, targeting the clustered hotspot mutations found in exon 8, with another pair of probes interrogating the E380Q mutation in exon 5. We assessed its sensitivity *in vitro* using synthetic oligonucleotides, harboring E380Q, L536R, Y537C, Y537N, Y537S or D538G mutations. Validation of the assay was performed on plasma samples from a prospective study and compared to next generation sequencing (NGS) data.

**Results:** The multiplex *ESR1*-ddPCR showed a high sensitivity with a limit of detection ranging from 0.07 to 0.19% in mutant allele frequency depending on the mutation tested. The screening of plasma samples from patients with AI-resistant metastatic breast cancer identified *ESR1* mutations in 29% of them with perfect concordance (and higher sensitivity) to NGS data obtained in parallel. Additionally, this test identifies patients harboring polyclonal alterations. Furthermore, the monitoring of ctDNA using this technique during treatment follow-up predicts the radiological response to palbociclib-fulvestrant.

**Conclusion:** The multiplex *ESR1*-ddPCR detects, in a single reaction, the most frequent *ESR1* activating mutations and is compatible with plasma samples. This method is thus suitable for real-time *ESR1* mutation monitoring in large cohorts of patients.

**Statement of translational relevance:** Exons 5 and 8 mutations in *ESR1* are recurrent mechanisms of resistance to aromatase inhibitors (AI) in estrogen receptor (ER)-positive metastatic breast cancer and may be targeted by selective ER down-regulators. We implemented a novel droplet digital PCR, which allows for the detection of the most frequent *ESR1* mutations in circulating cell-free DNA. In prospectively collected plasma samples, *ESR1* mutations were found in 29% of AI-resistant patients, with excellent concordance and higher sensitivity to next generation sequencing. Moreover, circulating *ESR1* mutations appear to be reliable markers for ctDNA monitoring in order to predict treatment response. Ultimately, the short turnaround time, high sensitivity and limited cost of the *ESR1*-ddPCR are compatible with repeated samplings to detect the onset of resistance to AI before the radiological progression. This opens a window of opportunity to develop new clinical strategies for breast cancer hormone therapy, as tested in an ongoing phase 3 trial.

**List of abbreviations:**

AI: Aromatase Inhibitor
cfDNA: Cell-free DNA
ctDNA: Circulating tumor DNA
ddPCR: Droplet digital PCR
ER+ HER2-MBC: ER+ HER2-negative Metastatic Breast Cancer
ER: Estrogen Receptor
*ER+*: Estrogen Receptor positive
LOB: Limit of blank
LOD: Limit of detection
MAF: Mutant Allele Frequency
PBMC: Peripheral blood mononuclear cells
PD: Progressive disease
SD: Standard deviation
ToP: Time of progression
WT: Wild type

**Human genes:**

*ESR1*: Estrogen Receptor 1
*HER2*: Human Epidermal Growth Factor Receptor 2
*EGFR*: Epithelial Growth Factor Receptor
*KRAS*: KRAS proto-oncogene, GTPase
*BRAF*: B-Raf Proto-Oncogene, Serine/Threonine kinase

## INTRODUCTION

In most breast cancer cases, the estrogen receptor (ER) exerts a key survival signaling, and ER-positive (ER+) immunostaining is a well-characterized predictive biomarker of hormone therapy efficiency at both early and advanced stages. In 2013, several studies reported activating point mutations in the estrogen receptor gene, *ESR1* (1–4). These oncogenic mutations activate the ER in a ligand-independent manner and were proposed to be a potential mechanism of resistance to hormone depletion, by aromatase inhibitors (AI) for example. In ER+ HER2-negative metastatic breast cancer (ER+ HER2-MBC) patients, *ESR1* mutations are rarely detected in primary tumors or at first metastatic relapse (5,6). However, their overall incidence increases alongside the exposure to AI (3–5,7), suggesting that *ESR1* mutations are a positively selected resistance mechanism to this treatment. It has been reported that 30 to 40% of ER+ HER2-MBC patients who are resistant to AI carry *ESR1* mutations, some even harboring polyclonal *ESR1* alterations (8,9). These mutations occur in the ligand-binding domain, typically in two hotspot locations: codon 380 (exon 5) and codons 536, 537 and 538 (exon 8) (10–12). Importantly, other hormone therapy agents, such as selective ER down-regulators (e.g., fulvestrant, elacestrant…), retain a significant activity on mutated ER (Supplementary Table 1) (12). This indicates that mutations in *ESR1* could be a predictive biomarker in the management of ER+ MBC (11). Accurate and sensitive detection tools are therefore needed to initiate clinical trials that investigate the clinical utility of *ESR1* mutation monitoring. In that context, a liquid biopsy approach based on circulating tumor DNA (ctDNA) detection is a most promising tool to detect the onset of resistance acquisition.

Our group, and others, have shown that drop-off droplet digital PCR (ddPCR) is an efficient tool to screen for clustered mutations that can be covered by a single TaqMan probe, unlike conventional ddPCR, which requires a specific probe for each targeted mutation. The drop-off assay relies on the concept that a single mismatch is sufficient to destabilize the complex ‘probe / target’ sequence. Hence, any mutation in the region covered by the ‘drop-off’ probe leads to a loss of signal when compared to wild-type (WT) samples. All previous studies reported high sensitivity and reproducibility for hotspot mutations in the *EGFR, KRAS*, and *BRAF* genes (13–15).

Here, we present the validation of a novel ddPCR assay, which combines a drop-off ddPCR, targeting the clustered hotspot mutations found in *ESR1* exon 8, with another pair of probes interrogating specifically the E380Q mutation, located in exon 5. This multiplex test detects, in a single reaction, up to 95% of the described activating *ESR1* mutations, based on previous studies (3,4,10,12,16), and is compatible with body fluid samples.

## MATERIALS AND METHODS

### Droplet digital PCR assay

#### TaqMan^®^ probes and primer design

For the E380Q mutation in exon 5, the assay used a probe specifically targeting the E380Q mutation (E380Q probe, FAM-labeled) and a reference probe (REFex5 probe, VIC-labeled) annealing to an adjacent invariant region (Figure 1A). For the clustered hotspot mutations in exon 8, a drop-off assay was designed using a probe targeting the WT sequence of codons 536, 537 and 538 where mutations are likely to be found (Hotspot probe, VIC-labeled) and a reference probe (REFex8 probe, FAM-labeled) annealing to an adjacent invariant region (Figure 1B). For compatibility with ctDNA detection, primers were designed to generate amplicons under 150 bp (101 bp and 125 bp for exons 5 and 8, respectively). The following primers were used to amplify (*a*) exon 5: Fwd primer: 5’-TTGCTTGTTTTCAGGCTTTGTGGA-3’; Rev primer 5’-AGCGCCAGACGAGACCAATCAT-3’; and (*b*) exon 8: Fwd primer: 5’-ACAGCATGAAGTGCAAGAACGT-3’; Rev primer 5’-TGGCTTTGGTCCGTCTCCTC-3’. The following TaqMan probes (Life Technologies) with a 5’ fluorophore and a 3’ non-fluorescent quencher (NFQ) were designed as follows: REFex5 5’-(VIC)-TGACCCTCCATGATC-(MGB NFQ)-3’; E380Q 5’-(6-FAM)-ACCTTCTACAATGTGCCTG-(MGB NFQ)-3’; REFex8 5’-(6-FAM)-CTAGCCGTGGAGGGGC-(MGB NFQ)-3’; Hotspot 5’-(VIC)-CCTCTATGACCTGCTGC-(MGB NFQ)-3’.

**Figure 1.**
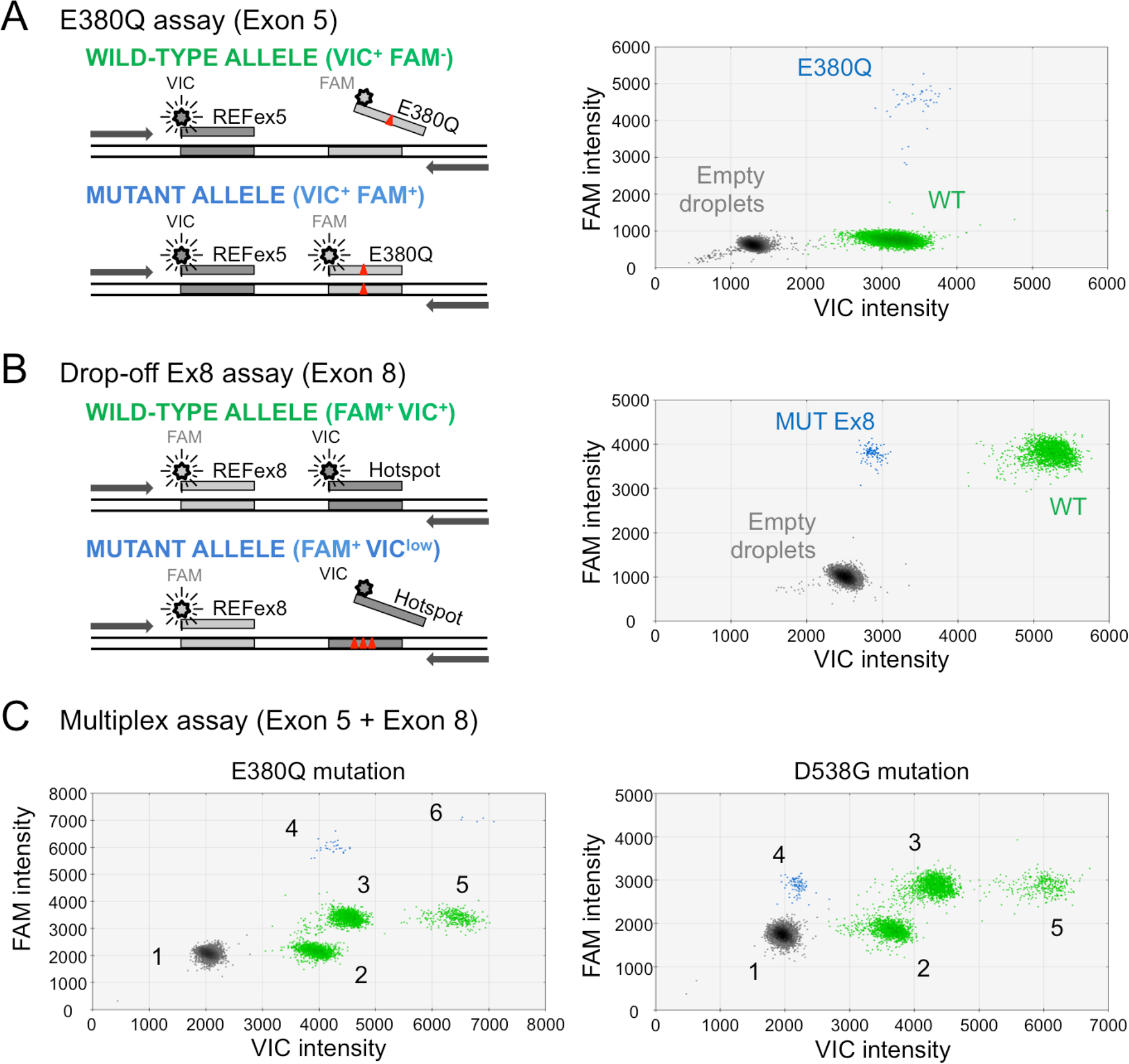
*ESR1* exons 5 and 8 assay design. **A.** Assay to detect the E380Q mutation in exon 5: wild-type alleles generate a signal from the reference probe only (VIC+, green cloud) whereas E380Q alleles generate a double positive signal (VIC+/FAM+, blue cloud). **B.** The Drop-off Ex8 assay detects the clustered hotspot mutations located in exon 8 (D538G in this example). Wild-type alleles generate a signal from both the reference and drop-off probes (FAM+/VIC+, green cloud) whereas alleles with alterations in codons 536-538 generate a signal from the reference probe only (FAM+, blue cloud). **C.** The multiplex assay detects, in a single reaction, the E380Q mutation (left panel) and mutations in exon 8 (right panel, D538G in this example). Cloud #1 = Empty droplets; Cloud #2 = WT Ex5; Cloud #3 = WT Ex8; Cloud #4 = E380Q (left panel) or MUT Ex8 (right panel); Cloud #5 = WT Ex5 + WT Ex8; Cloud #6 = E380Q + WT Ex5 + WT Ex8.

#### ddPCRmix preparation

ddPCR reactions were performed in 20 μL according to the manufacturer’s protocol. Briefly, 20 μL mastermix solution containing ddPCR™ Supermix for probes without dUTP (Bio-Rad Laboratories) at a final concentration of 1x, relevant primers at 450 nM each and relevant TaqMan^®^ probes (E380Q + REFex5 or Hotspot + REFex8 in simplex conditions or both assays combined in multiplex conditions) at 250 nM each (Applied Biosystems), DNA template (up to 8 μL) and nuclease-free water were loaded into a disposable droplet generator cartridge (Bio-Rad). After adding 70 μL of droplet generation oil (Bio-Rad), the cartridge was loaded into a QX100 Droplet Generator (Bio-Rad). Generated droplets were transferred to a 96-well PCR plate and PCR reactions were run on a C1000 thermal cycler (Bio-Rad) under the following program: 95°C 10 min, 40 cycles of (94°C 30 sec, 58°C 60 sec), 98°C 10 min. For optimization experiments, we used 20 ng (6,060 genome equivalent) of DNA per reaction. For the mutation screening in patient samples, we used 8 μL of cell-free DNA (cfDNA). Negative controls with no DNA and positive controls (E380Q and D538G mutations) with 6,060 WT genome-equivalent from peripheral blood mononucleated cells (PBMC) were included at each run. Reactions were analyzed on the Bio-Rad QX100 droplet reader.

#### ddPCR data analysis

The concentration of mutant DNA copies was estimated using the dedicated workflow available in the QuantaSoft v1.7.4 software. For the E380Q mutation, the number of mutant copies per droplet is equivalent to the number of VIC^+^/FAM^+^ droplets in simplex conditions and to clouds #4 and #6 in multiplex conditions (Figure 1). The mutant allele frequency (MAF) was calculated as follows: (VIC^+^/FAM^+^ droplets / (VIC^+^/FAM^+^ droplets + VIC^+^/FAM^−^ droplets)) for simplex conditions and (clouds #4 + #6/(clouds #4 + #6 + #2 + #5)) for multiplex conditions. For mutations in exon 8, the number of mutant copies per droplet is equivalent to the number of VIC^low^/FAM^+^ droplets in simplex conditions and to cloud #4 in multiplex conditions. The MAF was calculated as follows: (VIC^low^/FAM^+^ droplets / (VIC^low^/FAM^+^ droplets + VIC^+^/FAM^+^ droplets)) for simplex conditions and (cloud #4 /(clouds #4 + #3 + #5)) for multiplex conditions. Samples were run in triplicates and were considered to be positive if the merged replicates presented a minimum of 3 mutant droplets for E380Q or 8 mutant droplets for exon 8 and if the average MAF was higher than the defined LOD of the multiplex assay.

### *In vitro* performance

The limit of blank (LOB) was determined as previously reported (13,15,17,18). Briefly, we defined the false-positive mean and associated standard deviation (SD) of the E380Q and Drop-off Ex8 assays, in simplex or multiplex conditions by analyzing 48 replicates of WT genomic DNA extracted from PBMC. Then, the calculated 95% confidence interval was used to define the LOB. To assess the limit of detection (LOD) of the E380Q and Drop-off Ex8 assays, we used synthetic oligonucleotides harboring E380Q, or the most frequent mutations in exon 8 (L536R, Y537C, Y537N, Y537S or D538G). Serial dilutions in 10 ng (3,030 genome equivalent) of WT DNA from PBMC, reproducing MAFs from 0.8% to 0.04%, were analyzed in 8 replicates for each mutated oligonucleotide.

### Validation in clinical samples

All plasma samples analyzed for *ESR1* mutation status were collected from patients with AI-resistant ER+ HER2-MBC. Patients were prospectively enrolled at the Institut Curie (Paris, France) in the ethically-approved ALCINA study (NCT02866149, cohort 6) after signed informed consent and that were included prior to the initiation of a new line of therapy with palbociclib and fulvestrant. Progression-free survival (PFS), defined as the time from inclusion in the study to progression disease (PD) or death from any cause, was collected prospectively. Survival analysis was performed using Kaplan–Meier plots with significance tested using the log-rank test. Plasma was isolated from fresh blood collected in EDTA blood collection tubes (BD Vacutainer^®^) within 3 hours as previously performed in the laboratory (19,20) and stored at −80°C until needed. For *ESR1* screening, 2 mL of plasma were thawed and cfDNA extracted using the QIAamp^®^ Circulating Nucleic Acid Kit (Qiagen) according to the manufacturer’s protocol and stored at −20°C until use. In parallel to ddPCR experiments, targeted-NGS on a panel of 39 cancer-related genes was performed in a blind fashion to allow head-to-head comparison with the *ESR1*-ddPCR assay. Following library preparation, samples were subjected to ultra-deep sequencing on Illumina HiSeq2500 using a 2 × 100 bp paired-end configuration. Read depth obtained for the two *ESR1* hotspots regions was higher than 5,800X. Paired-end read alignments were performed on GRCh37 (hg19) human reference with Bowtie2 (v2.1.0). Once aligned, paired-end reads that map to multiple locations or with poor mapping quality (score <6) were removed. Pileup files were generated using samtools (v.1.1) and variant calling was performed using Varscan2 (v2.3.6). A minimum base quality of 15 was required to count for a read at a position, and only variants supported by a minimum of 5 mutated reads at a position with a minimum read depth of 12 were selected. Additionally, the nucleotide composition for each position of the regions containing the two *ESR1* hotspots was extracted from the BAM alignment files using GATK (v.3.5).

## RESULTS

### Screening of multiple *ESR1* hotspot mutations by ddPCR

Based on the evidence that more than 80% of the activating *ESR1* mutations found in ER+ HER2-MBC patients resistant to AI alters codon 380 in exon 5 and the codons 536, 537 and 538 in exon 8 (19% and 64% respectively, see Supplementary Table 1), we developed a ddPCR assay targeting these two regions. This assay is composed of only two pairs of TaqMan probes, which can be combined in a single ddPCR reaction. The first pair targets specifically the E380Q mutation in exon 5, using a probe complementary to the E380Q mutant allele (E380Q probe, FAM-labeled) and a reference probe spanning an adjacent invariable region (REFex5 probe, VIC-labeled) (Figure 1A, left panel). In this unconventional E380Q assay, droplets containing WT alleles are VIC-positive only, because the E380Q probe cannot hybridize onto WT sequences, whereas droplets containing E380Q mutant alleles are double positive (VIC^+^/FAM^+^) (Figure 1A, right panel). The second pair of probes, constituting the Drop-off Ex8 assay, targets the clustered mutations in exon 8 using a probe complementary to the WT sequence of the altered region (Hotspot probe, VIC-labeled), which detects all the mutations occurring at codons 536, 537 and 538, together with a reference probe spanning an invariable region within the same amplicon (REFex8 probe, FAM-labeled) (Figure 1B, left panel). Therefore, droplets containing WT alleles are double positive (FAM^+^/VIC^+^) whereas the mismatch induced by the mutations leads to a VIC signal decrease (FAM^+^/VIC^low^), resulting in a shift of the droplet cloud (MUT Ex8) toward a single FAM-positive population (Figure 1B, right panel).

The multiplexed assay, combining primers and probes from the E380Q and the Drop-off Ex8 assays, allows to screen for mutations in exons 5 and 8 in a single reaction. In a test using synthetic E380Q or D538G oligonucleotides, we could distinguish each WT and mutant clouds for exons 5 and 8 in multiplex conditions (Figure 1C). We observed an additional cloud of droplets containing both WT Ex5 and WT Ex8 amplicons (Figure 1C, Cloud #5), which we also find in WT samples (data not shown), as well as a cloud containing E380Q and WT copies when testing for the E380Q mutation (Figure 1C, left panel, Cloud #6), and not observed when testing for the D538G mutation.

### Detection of polyclonal alterations

In addition to the D538G mutation, we tested four other synthetic oligonucleotides harboring the Y537C, Y537N, Y537C or L536R mutations found in exon 8. We observed that the position of the MUT Ex8 cloud was dependent on the mutation tested (Figure 2A). This suggests that each nucleotide change does not equally destabilize the complex ‘probe / target sequence’. The largest shifts were observed for the mutations D538G and Y537C. Both clouds of droplets containing these mutant alleles were localized in the same area and were therefore indistinguishable from each other; whereas the clouds for Y537N, Y537S and L536R displayed smaller shifts and were distinct from the D538G/Y537C cloud. We further tested combinations of two mutant oligonucleotides in multiplex conditions confirming that multiple mutations in exon 8 (e.g., D538G + Y537N, D538G + Y537S, D538G + L536R or Y357N + L536R) display a specific pattern (Figure 2B). We clearly detected the combination of one mutation in exon 8 (e.g., D538G) with E380Q (Figure 2B). The multiplex *ESR1*-ddPCR can thus identify patients carrying polyclonal alterations, as previously reported (8, 9).

**Figure 2.**
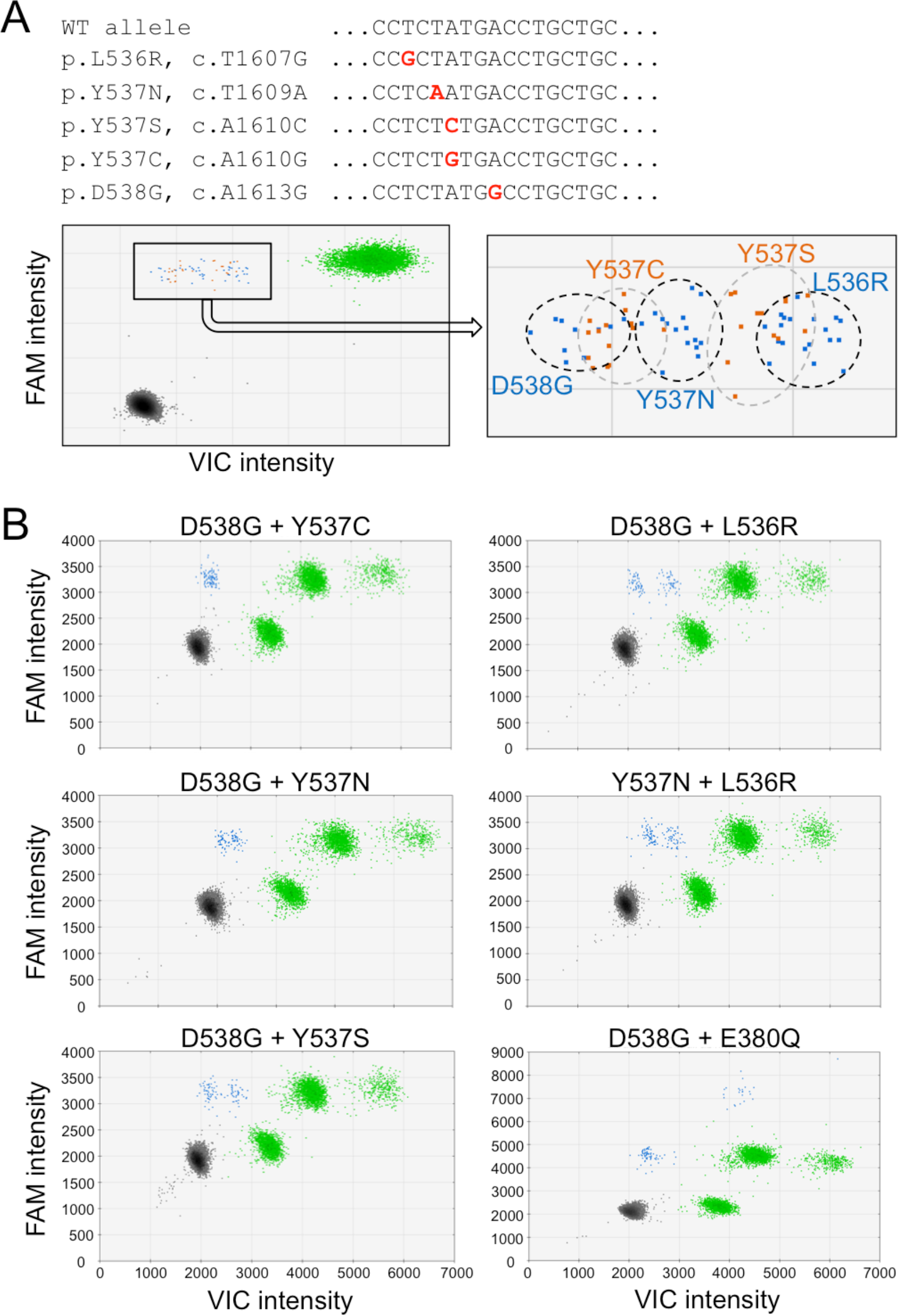
Detection of *ESR1* polyclonal mutations. **A.** Most frequent mutations located in *ESR1* exon 8 targeted by the Drop-off Ex8 assay. **B.** Representative examples of double mutations (synthetic oligonucleotides combined) detected in multiplex condition.

### *In vitro* performances

We further estimated the specificity and sensitivity of the *ESR1*-ddPCR by analyzing 48 replicates of pure WT DNA and serial dilutions of the mutant synthetic oligonucleotides recapitulating MAFs from 0.8% to 0.04% (see Methods section for more details). The *ESR1*-ddPCR assay showed high specificity with a maximum of one false-positive event observed per reaction (Figure 3A and Supplementary Table 2). The limit of blank (LOB) was estimated at 0.004% for exon 5 mutations and 0.008% for exon 8 mutations. The limit of detection (LOD), defined as the lowest MAF with all replicates presenting values above the LOB, was estimated at 0.19% in mutant allele frequency for E380Q (Figure 3B and Supplementary Table 1) and ranged from 0.07 to 0.13% depending on the exon 8 mutation tested (Figure 3C and Supplementary Table 1). LOB and LOD were also tested in simplex conditions and similar values were observed (LOB were 0.003% for the E380Q assay and 0.01% for the Drop-off Ex8 assay; LOD were 0.09% for mutation E380Q and ranging from 0.06 to 0.1% depending on the exon 8 mutation tested, see Supplementary Figure 1A).

**Figure 3.**
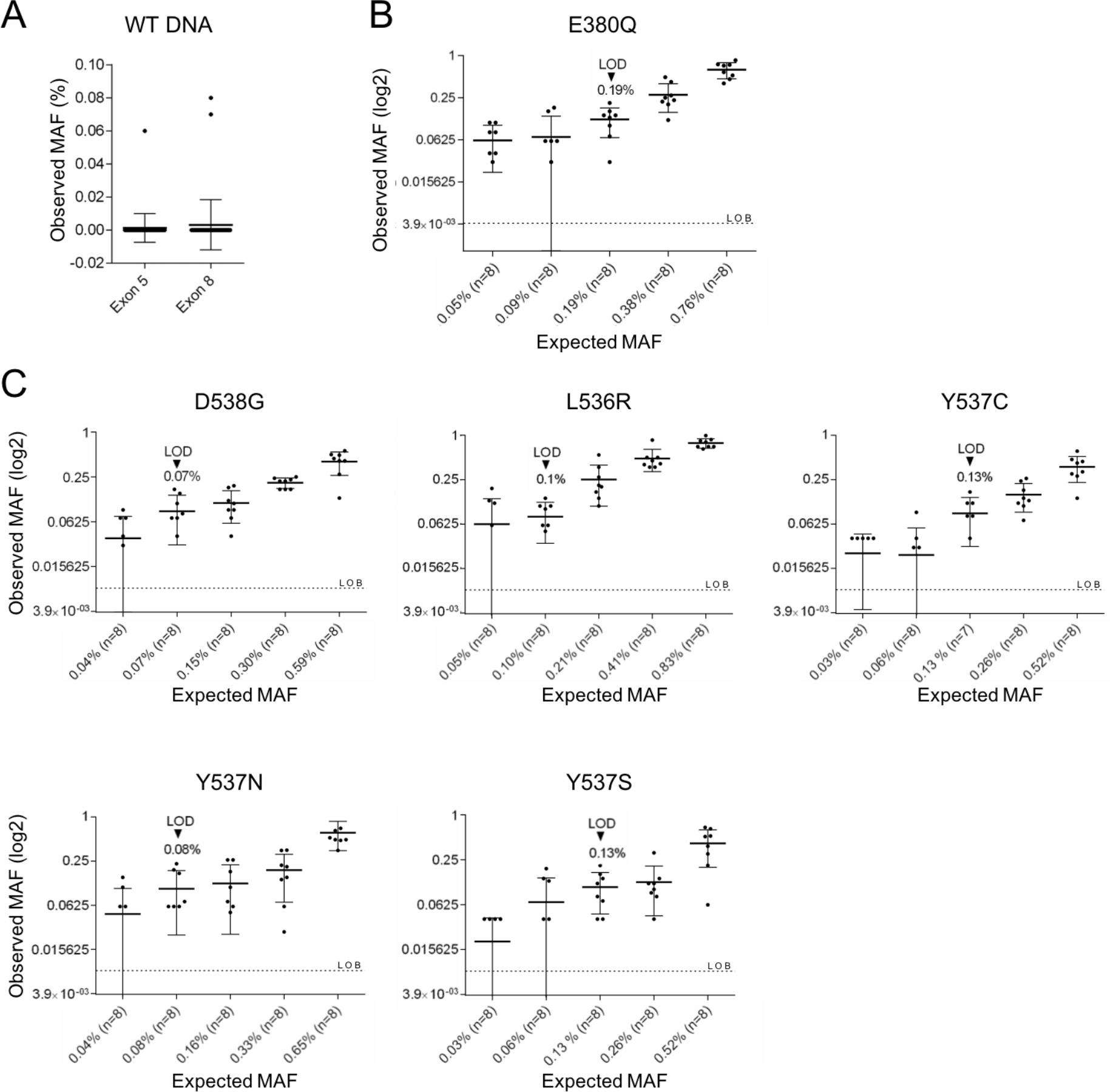
*In vitro* performance of the multiplex *ESR1*-ddPCR. **A.** False positive events for exons 5 or 8 mutations observed from pure WT DNA tested with the multiplex *ESR1*-ddPCR. **B.** LOD estimation for exon 5 E380Q mutation. **C.** LOD estimation for exon 8 mutations D538G, L536R, Y537C, Y537N or Y537S. See method section for more details and Supplementary Table 1. LOB: limit of blank, LOD: limit of detection, estimated as the 95% CI of the mean false-positive calls.

### Validation in clinical samples

To validate the performance of the *ESR1* multiplex ddPCR assay, we tested a series of 43 plasma samples from a prospective cohort of patients with HR+ HER2-MBC progressing under hormone therapy. We successfully detected *ESR1* mutations in 11 out of the 42 (26%) informative patient samples (Table 1). Four cases (P-05, P-17, P-37 and P-43) harbored an E380Q mutation (36% of the mutant cases, Figure 4A) and 8 cases (P-08, P-18, P-20, P-25, P-28, P35, P-39 and P-43) carried at least one mutation in exon 8 (73% of the mutant cases, Figure 4B). Interestingly, based on the shape and the location of the MUT Ex8 clouds, we detected that samples P-18 and P-28 harbored multiple mutations (Figure 4C). We also observed that P-43 harbored *ESR1* mutations in both exons 5 and 8 (Figure 4C). In parallel, targeted-NGS, including the *ESR1* gene, was performed on 32 plasma samples, for which we had sufficient cfDNA, to allow head-to-head comparison with the *ESR1*-ddPCR assay. Importantly, NGS analysis was performed in a blinded fashion, with no prior knowledge of ddPCR results. The samples sequenced included 10 mutant and 22 WT cases according to ddPCR analysis. All *ESR1* mutations detected by ddPCR were confirmed by NGS except for 2 cases (P-17 and P-20), in which the *ESR1* MAF was found to be <1% by ddPCR (Table 1, Supplementary Figure 2A). P-25 was not tested with NGS due to lack of cfDNA (Table 1, Supplementary Figure 2B). We also confirmed the WT status of the 22 remaining plasma samples. In case P-18, NGS results indicated that the mutations were D538G and L536H, a combination we did not test before (Table 1). In case P-28, we observed 3 mutations in exon 8: L536H, Y537N and Y537S. Interestingly, for case P-35, NGS showed that the mutation in exon 8 was Y537N, which did not result from the usual C>T substitution in position 1609, but from a small deletion/insertion at nucleotides 1608-1609 (c.1608_1609delinsTA). Overall, the *ESR1* multiplex ddPCR performs better than standard targeted NGS in identifying *ESR1* mutation carriers owing to its improved sensitivity.

**Table 1:**
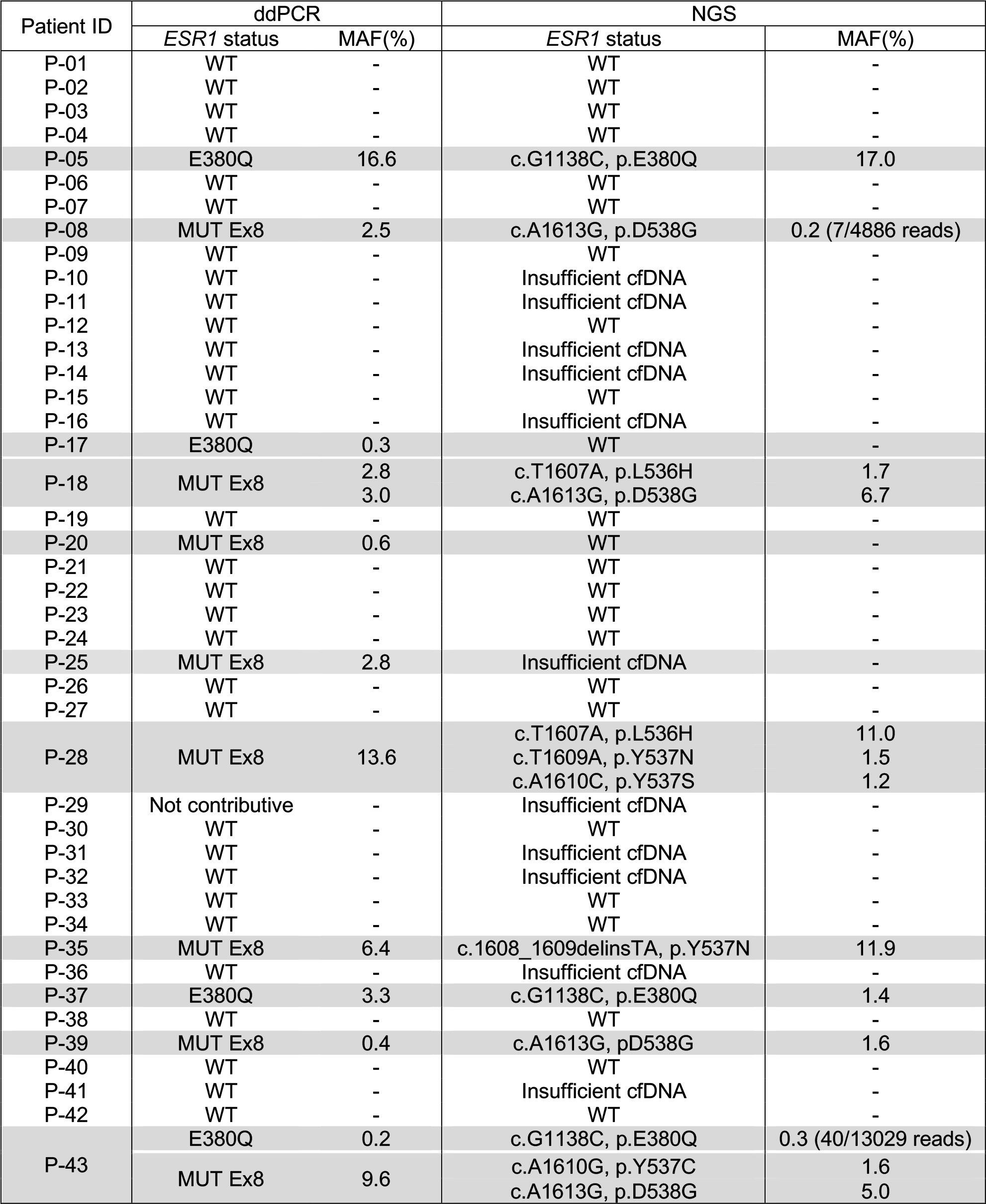
Detection of *ESR1* mutations using ddPCR and NGS in plasma samples from patients with MBC. Samples with mutation identified by ddPCR are highlighted in grey. Corresponding ddPCR profiles are presented in Figure 4 and Supplemental Figure 2.

**Figure 4.**
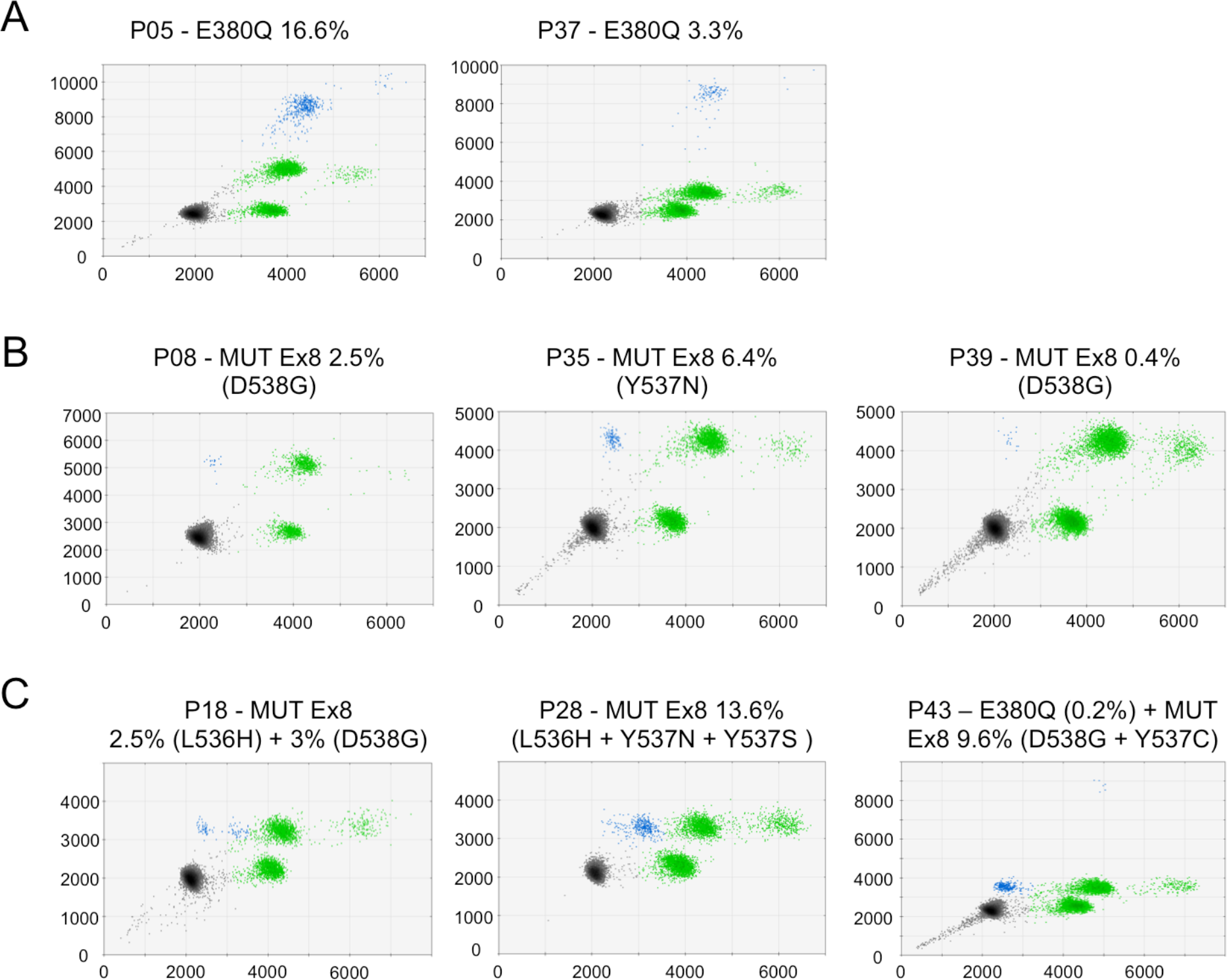
Validation of the *ESR1* ddPCR multiplex assay on patient plasma samples. Representative ddPCR profiles for E380Q mutation **(A)**, exon 8 mutations (D538G or Y357N) **(B)** and polyclonal *ESR1* mutations **(C)**. Case P-18, double exon 8 mutations identified as L536H and D538G; case P-28, exon 8 mutations identified as L536H, Y537N and Y537S and case P-43, exon 5 + exon 8 mutations identified as D538G, Y537C and E380Q by NGS. FAM and VIC intensities are shown in arbitrary units. MAFs determined by ddPCR are indicated above each plot together with the mutation identified by NGS in brackets.

### Monitoring of circulating *ESR1* mutant copies to predict response to palbociclib-fulvestrant therapy

We next analyzed the impact of the *ESR1* mutant status, detected with the *ESR1*-ddPCR, in plasma samples collected at baseline and during treatment follow-up. To perform this analysis, we extended the cohort to 60 patients. For each patient, four blood samples were collected: before treatment (D0), after 15 days (D15) and 30 days (D30) of treatment and at the time of progression (ToP). Among the 59 patients screened with contributive results, *ESR1* mutations were detected in 17 (28.8%), which is in line with proportions of patients progressing under AI treatment previously reported (8, 9, 11, 24). Out of the 17 patients carrying an *ESR1* mutation, 15 had an evaluable PFS (2 patients were withdrawn from the study shortly after the treatment initiation, Figure 5A). First, we observed no impact of the *ESR1* mutational status (mutant vs wild-type) detected at baseline on the PFS (Figure S3A). This is in agreement with previous data from the PALOMA-3 trial reporting no improved PFS for patients carrying *ESR1* mutations at baseline and treated with fulvestrant (11, 24). After 3 months of treatment, 6 patients presented a disease progression according to RECIST criteria (Figure 5A). We further analyzed the dynamics of *ESR1* mutant copy levels in regards to the 3-months disease progression status (PD versus non-PD). At baseline, we observed a median number of ctDNA copies of 54 per ml of plasma (mean = 188.5, range [3 - 1233]) and there was no apparent association with disease progression at 3 months (Figure 5B). After 15 days of treatment, we uncovered a dramatic decrease (median = 0, mean = 14.3, range [0 - 74]) with 58% (7/12) of patients reaching undetectable *ESR1* mutant levels (Figure 5B). The evolution of the number of ctDNA copies between D0 and D15, for each patient, demonstrates a decrease for all (Figure 5C, Supplementary Table 3). To a minor extent, a significant decrease was also observed on total cfDNA at D15 (Figure S3B-C), which was previously described and related to the cytostatic growth arrest triggered by palbociclib (24). At D30, the majority of patients with early disease progression had increased ctDNA levels whereas the majority of patients with longer PFS displayed decreased or stable ctDNA levels (Figure 5C). Overall, a lower proportion of patients displayed undetectable ctDNA levels (53% = 8/15, Figure 5B), the majority of which (87.5% = 7/8) had no early disease progression. Patients with early disease progression had significantly more ctDNA detected at D30 than patients with longer PFS (p < 0.0001; Figure 5D). Only one out of 6 patients with early disease progression had no ctDNA detected whereas only 2 out of 9 patients without early disease progression showed detectable ctDNA (Figure 5B). This ctDNA positivity at D30 correlates with PFS (p = 0.0063; Figure 5E-F). Here, we show that the detection of ctDNA after 30 days of palbociclib-fulvestrant is a promising dynamic biomarker associated with PFS.

**Figure 5.**
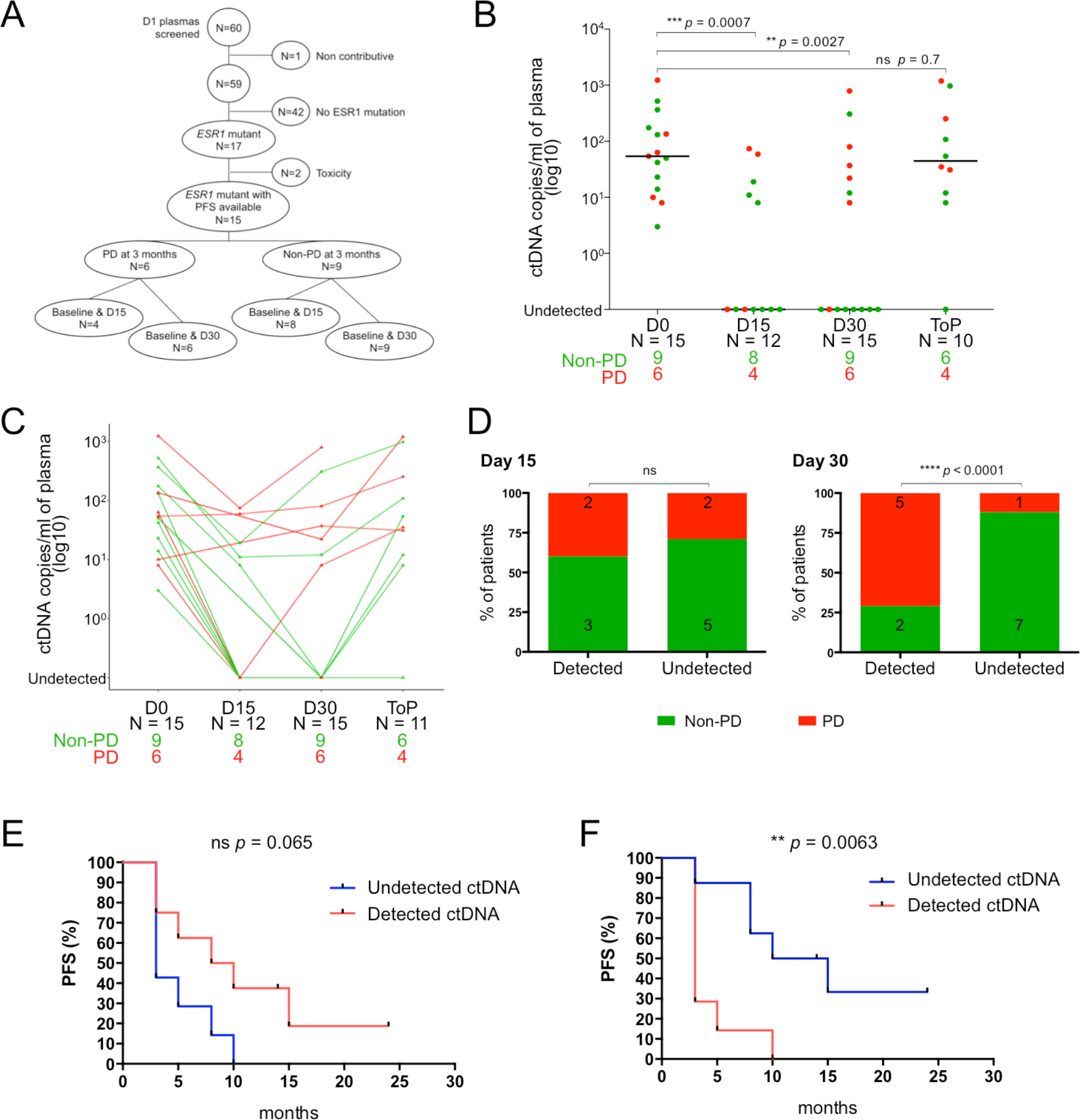
Monitoring of circulating *ESR1* mutant copies to predict response to palbociclib-fulvestrant therapy. **A.** Flow chart of plasma samples analyzed from the ALCINA cohort 6 ‘Palbociclib’ (NCT02866149) for *ESR1* mutations with the *ESR1*-ddPCR multiplex. PD: progressive disease, Non-PD: regressive or stable disease. **B.** Number of ctDNA copies detected per ml of plasma collected at the 4 time points during treatment follow-up (D0, N = 15; D15, N = 12; D30, N = 15; ToP, N = 10). **C.** ctDNA dynamics observed during treatment follow-up in patients experiencing progressive disease (PD, red) versus regressive or stable disease (non-PD, green) at 3 months according to RECIST criteria. (D0: non-PD = 9, PD = 6; D15: non-PD = 8, PD = 4; D30: non-PD = 9, PD = 6; ToP: non-PD = 6, PD = 4). **D.** Two sample test proportions for detectable and undetectable ctDNA distribution at D15 and D30. **E, F.** Survival (PFS) according to ctDNA detection at D15 **(E)** sor D30 **(F)**.

## DISCUSSION

We successfully developed a ddPCR assay detecting the most frequent activating *ESR1* mutations at once that is compatible with liquid biopsies. By using an unconventional design, which includes a drop-off assay, we targeted, in a single reaction, the E380Q mutation and all the mutations occurring at codons 536 to 540. The multiplex *ESR1*-ddPCR covers >80% of the currently described *ESR1* mutations and >90% of functionally characterized activating *ESR1* mutations. Several teams have previously developed ddPCR assays which target only the most frequent *ESR1* mutations found in exon 8: D538G, Y537S, Y537N and Y537C (6,21–23). These assays were designed following the conventional ddPCR method containing specific TaqMan probes complementary to each mutant or WT allele. This implies that each mutation is tested in a separate reaction. Thus, multiplex assays were developed to reduce the number of reactions (9, 11). However, these assays cannot identify more than 4 *ESR1* mutations in a single reaction and the mutant samples were usually confirmed by singleplex tests (24). To our knowledge, we are the first to have developed a ddPCR assay which can detect, in a single reaction, at least eight different mutations in *ESR1*, namely: E380Q, L536H, L536R, Y537C, Y537N (T>A), Y537N (delinsTA), Y537S and D538G. In addition, our system can identify samples harboring multiple *ESR1* mutations (e.g., E380Q combined with one or more mutations in exon 8). Polyclonal *ESR1* mutations are well-described events (9,10) and the *ESR1*-ddPCR assay would be useful in monitoring the dynamics of each mutation during treatment follow up as seen for P-43. The multiplex *ESR1*-ddPCR assay is highly sensitive, detecting all tested mutations at frequencies lower than 0.19%, an improvement as compared with NGS. We also demonstrated that the *ESR1*-ddPCR is highly specific by cross-validation with NGS experiments. Lupini *et al.* recently developed an assay based on an ‘enhanced-ice-COLD-PCR followed by NGS’ with a sensitivity reaching 0.01% (25). However, this ddPCR assay targets specifically the Y537S mutation and involves an enrichment step of the mutant copies preceding the ddPCR assay. Yet, in a context of patient monitoring by liquid biopsy, biological samples are of limited quantity and must be tested rapidly at a low cost. The multiplex *ESR1*-ddPCR can detect most *ESR1* mutations in a single reaction faster and at a lower cost than NGS or any other currently available technology.

Interestingly, we observed that exons 5 and 8 mutations can be easily distinguished. Moreover, any nucleotide change covered by the Drop-off Ex8 assay can be identified, as confirmed by the detection of the previously unreported mutation Y537N (delinsTA). We observed that, among exon 8 mutations, the shift in clouds is unique depending on the mutation, indicating if the mutation is more likely to be a D538G or Y537C allele versus mutations in codon 536 or other changes in codon 537. However, the fact that this assay does not recover the exact nucleotide change in exon 8 constitutes its most prominent limitation. From a clinical perspective, all mutations in codons 536-538 have been equally associated with resistance to AI therapy. Nonetheless, preclinical data suggest that the Y537S mutation, which accounts for about 10% of all *ESR1* mutations, may be less sensitive to fulvestrant than other mutations (12). If this observation is confirmed to be clinically relevant, the *ESR1*-ddPCR could be used as a first screening tool, since the shift associated to Y537S is distinguishable from the most frequent mutation: D538G, followed by subsequent sequencing of exon 8, to distinguish Y537S from other 536/537 mutations. A second theoretical limitation is that synonymous mutations may also be detected by this assay. However, *ESR1* synonymous mutations, as well as polymorphisms in codons 536-540, are extremely rare. According to the Exome Aggregation Consortium, only two synonymous mutations, L536L and D538D were described at very low frequencies, 6.303e^−05^ and 8.251e^−06^, respectively.

The improved analytical sensitivity of the *ESR1*-ddPCR is particularly useful to monitor ctDNA during treatment follow-up. We demonstrated that *ESR1* mutations are good markers for ctDNA dynamics exploration and prediction of treatment response. Indeed, we observed that detection of ctDNA after 30 days of palbociclib-fulvestrant, using the *ESR1*-ddPCR, correlates with the treatment response and has an impact on PFS.

In conclusion, this method presents the advantage to screen for at least 80% of the *ESR1* mutations in a single reaction, as required by large screening studies involving plasma samples. This assay is currently implemented in the randomized phase 3 PADA-1 trial (NCT03079011) in which it allows for real-time monitoring of *ESR1* mutation rising in 1000 ER+ HER2-MBC patients treated with palbociclib and AI.

## ACKNOWLEDGEMENTS

We would like to thank members of the Circulating Tumor Biomarkers laboratory for helpful discussions and especially Amanda Silveira for critical reading of the manuscript. High-throughput sequencing has been performed by the ICGex NGS platform of the Institut Curie supported by the grants ANR-10-EQPX-03 (Equipex) and ANR-10-INBS-09-08 (France Génomique Consortium) from the Agence Nationale de la Recherche (“Investissements d’Avenir” program), by the Canceropole Ile-de-France and by the SiRIC-Curie program - SiRIC Grant « INCa-DGOS-Inserm_12554 ».

## REFERENCES

1. Li S, Shen D, Shao J, Crowder R, Liu W, Prat A, et al. Endocrine-therapy-resistant ESR1 variants revealed by genomic characterization of breast-cancer-derived xenografts. Cell Rep;4(6):1116–30 doi S2211-1247(13)00463-4[pii]10.1016/j.celrep.2013.08.022.

2. Merenbakh-Lamin K, Ben-Baruch N, Yeheskel A, Dvir A, Soussan-Gutman L, Jeselsohn R, et al. D538G mutation in estrogen receptor-alpha: A novel mechanism for acquired endocrine resistance in breast cancer. Cancer Res;73(23):6856–64 doi 0008-135472.CAN-13-1197[pii]1410.1158/0008-5472.CAN-13-1197.

3. Robinson DR, Wu YM, Vats P, Su F, Lonigro RJ, Cao X, et al. Activating ESR1 mutations in hormone-resistant metastatic breast cancer. Nat Genet;45(12):1446–51 doi ng.2823[pii]10.1038/ng.2823.

4. Toy W, Shen Y, Won H, Green B, Sakr RA, Will M, et al. ESR1 ligand-binding domain mutations in hormone-resistant breast cancer. Nat Genet;45(12):1439–45 doi ng.2822[pii]10.1038/ng.2822.

5. Jeselsohn R, Yelensky R, Buchwalter G, Frampton G, Meric-Bernstam F, Gonzalez-Angulo AM, et al. Emergence of constitutively active estrogen receptor-alpha mutations in pretreated advanced estrogen receptor-positive breast cancer. Clin Cancer Res;20(7):1757–67 doi 10.1158/1078-0432.CCR-13-2332.

6. Takeshita T, Yamamoto Y, Yamamoto-Ibusuki M, Inao T, Sueta A, Fujiwara S, et al. Droplet digital polymerase chain reaction assay for screening of ESR1 mutations in 325 breast cancer specimens. Transl Res;166(6):540–53 e2 doi S1931-5244(15)00306-0[pii]10.1016/j.trsl.2015.09.003.

7. Schiavon G, Hrebien S, Garcia-Murillas I, Cutts RJ, Pearson A, Tarazona N, et al. Analysis of ESR1 mutation in circulating tumor DNA demonstrates evolution during therapy for metastatic breast cancer. Sci Transl Med;7(313):313ra182 doi 7/313/313ra182[pii]10.1126/scitranslmed.aac7551.

8. Chandarlapaty S, Chen D, He W, Sung P, Samoila A, You D, et al. Prevalence of ESR1 Mutations in Cell-Free DNA and Outcomes in Metastatic Breast Cancer: A Secondary Analysis of the BOLERO-2 Clinical Trial. JAMA Oncol;2(10):1310–5 doi 2542919[pii]10.1001/jamaoncol.2016.1279.

9. Takeshita T, Yamamoto Y, Yamamoto-Ibusuki M, Tomiguchi M, Sueta A, Murakami K, et al. Analysis of ESR1 and PIK3CA mutations in plasma cell-free DNA from ER-positive breast cancer patients. Oncotarget;8(32):52142–55 doi 10.18632/oncotarget.1847918479[pii].

10. Chung JH, Pavlick D, Hartmaier R, Schrock AB, Young L, Forcier B, et al. Hybrid capture-based genomic profiling of circulating tumor DNA from patients with estrogen receptor-positive metastatic breast cancer. Ann Oncol;28(11):2866–73 doi 4098869[pii]10.1093/annonc/mdx490.

11. Fribbens C, O’Leary B, Kilburn L, Hrebien S, Garcia-Murillas I, Beaney M, et al. Plasma ESR1 Mutations and the Treatment of Estrogen Receptor-Positive Advanced Breast Cancer. J Clin Oncol;34(25):2961–8 doi JCO.2016.67.3061[pii]10.1200/JCO.2016.67.3061.

12. Toy W, Weir H, Razavi P, Lawson M, Goeppert AU, Mazzola AM, et al. Activating ESR1 Mutations Differentially Affect the Efficacy of ER Antagonists. Cancer Discov;7(3):277–87 doi 2159-8290.CD-15-1523[pii]10.1158/2159-8290.CD-15-1523.

13. Decraene C, Silveira AB, Bidard FC, Vallee A, Michel M, Melaabi S, et al. Multiple Hotspot Mutations Scanning by Single Droplet Digital PCR. Clin Chem;64(2):317–28 doi clinchem.2017.272518[pii]10.1373/clinchem.2017.272518.

14. Seki Y, Fujiwara Y, Kohno T, Takai E, Sunami K, Goto Y, et al. Picoliter-Droplet Digital Polymerase Chain Reaction-Based Analysis of Cell-Free Plasma DNA to Assess EGFR Mutations in Lung Adenocarcinoma That Confer Resistance to Tyrosine-Kinase Inhibitors. Oncologist;21(2):156–64 doi theoncologist.2015-0288[pii]10.1634/theoncologist.2015-0288.

15. Bidshahri R, Attali D, Fakhfakh K, McNeil K, Karsan A, Won JR, et al. Quantitative Detection and Resolution of BRAF V600 Status in Colorectal Cancer Using Droplet Digital PCR and a Novel Wild-Type Negative Assay. J Mol Diagn;18(2):190–204 doi S1525-1578(15)00262-7[pii]10.1016/j.jmoldx.2015.09.003.

16. Niu J, Andres G, Kramer K, Kundranda MN, Alvarez RH, Klimant E, et al. Incidence and clinical significance of ESR1 mutations in heavily pretreated metastatic breast cancer patients. Onco Targets Ther;8:3323–8 doi 10.2147/OTT.S92443ott-8-3323[pii].

17. Zonta E, Garlan F, Pecuchet N, Perez-Toralla K, Caen O, Milbury C, et al. Multiplex Detection of Rare Mutations by Picoliter Droplet Based Digital PCR: Sensitivity and Specificity Considerations. PLoS One;11(7):e0159094 doi 10.1371/journal.pone.0159094PONE-D-16-13557[pii].

18. Milbury CA, Zhong Q, Lin J, Williams M, Olson J, Link DR, et al. Determining lower limits of detection of digital PCR assays for cancer-related gene mutations. Biomol Detect Quantif;1(1):8–22 doi 10.1016/j.bdq.2014.08.001S2214-7535(14)00004-7[pii].

19. Madic J, Kiialainen A, Bidard FC, Birzele F, Ramey G, Leroy Q, et al. Circulating tumor DNA and circulating tumor cells in metastatic triple negative breast cancer patients. Int J Cancer;136(9):2158–65 doi 10.1002/ijc.29265.

20. Lebofsky R, Decraene C, Bernard V, Kamal M, Blin A, Leroy Q, et al. Circulating tumor DNA as a non-invasive substitute to metastasis biopsy for tumor genotyping and personalized medicine in a prospective trial across all tumor types. Mol Oncol;9(4):783-90 doi S1574-7891(14)00288-9[pii]10.1016/j.molonc.2014.12.003.

21. Clatot F, Perdrix A, Augusto L, Beaussire L, Delacour J, Calbrix C, et al. Kinetics, prognostic and predictive values of ESR1 circulating mutations in metastatic breast cancer patients progressing on aromatase inhibitor. Oncotarget;7(46):74448–59 doi 12950[pii]10.18632/oncotarget.12950.

22. Gyanchandani R, Kota KJ, Jonnalagadda AR, Minteer T, Knapick BA, Oesterreich S, et al. Detection of ESR1 mutations in circulating cell-free DNA from patients with metastatic breast cancer treated with palbociclib and letrozole. Oncotarget;8(40):66901–11 doi 10.18632/oncotarget.1138311383[pii].

23. Wang P, Bahreini A, Gyanchandani R, Lucas PC, Hartmaier RJ, Watters RJ, et al. Sensitive Detection of Mono- and Polyclonal ESR1 Mutations in Primary Tumors, Metastatic Lesions, and Cell-Free DNA of Breast Cancer Patients. Clin Cancer Res;22(5):1130–7 doi 1078-0432.CCR-15-1534[pii]10.1158/1078-0432.CCR-15-1534.

24. O’Leary B, Hrebien S, Morden JP, Beaney M, Fribbens C, Huang X, et al. Early circulating tumor DNA dynamics and clonal selection with palbociclib and fulvestrant for breast cancer. Nat Commun;9(1):896 doi 10.1038/s41467-018-03215-x10.1038/s41467-018-03215-x[pii].

25. Lupini L, Moretti A, Bassi C, Schirone A, Pedriali M, Querzoli P, et al. High-sensitivity assay for monitoring ESR1 mutations in circulating cell-free DNA of breast cancer patients receiving endocrine therapy. Sci Rep;8(1):4371 doi 10.1038/s41598-018-22312-x10.1038/s41598-018-22312-x[pii].

