## Supplementary figures for "A single droplet digital PCR for *ESR1* activating mutations detection in plasma"

### Supplementary materials

### Supplementary Figures

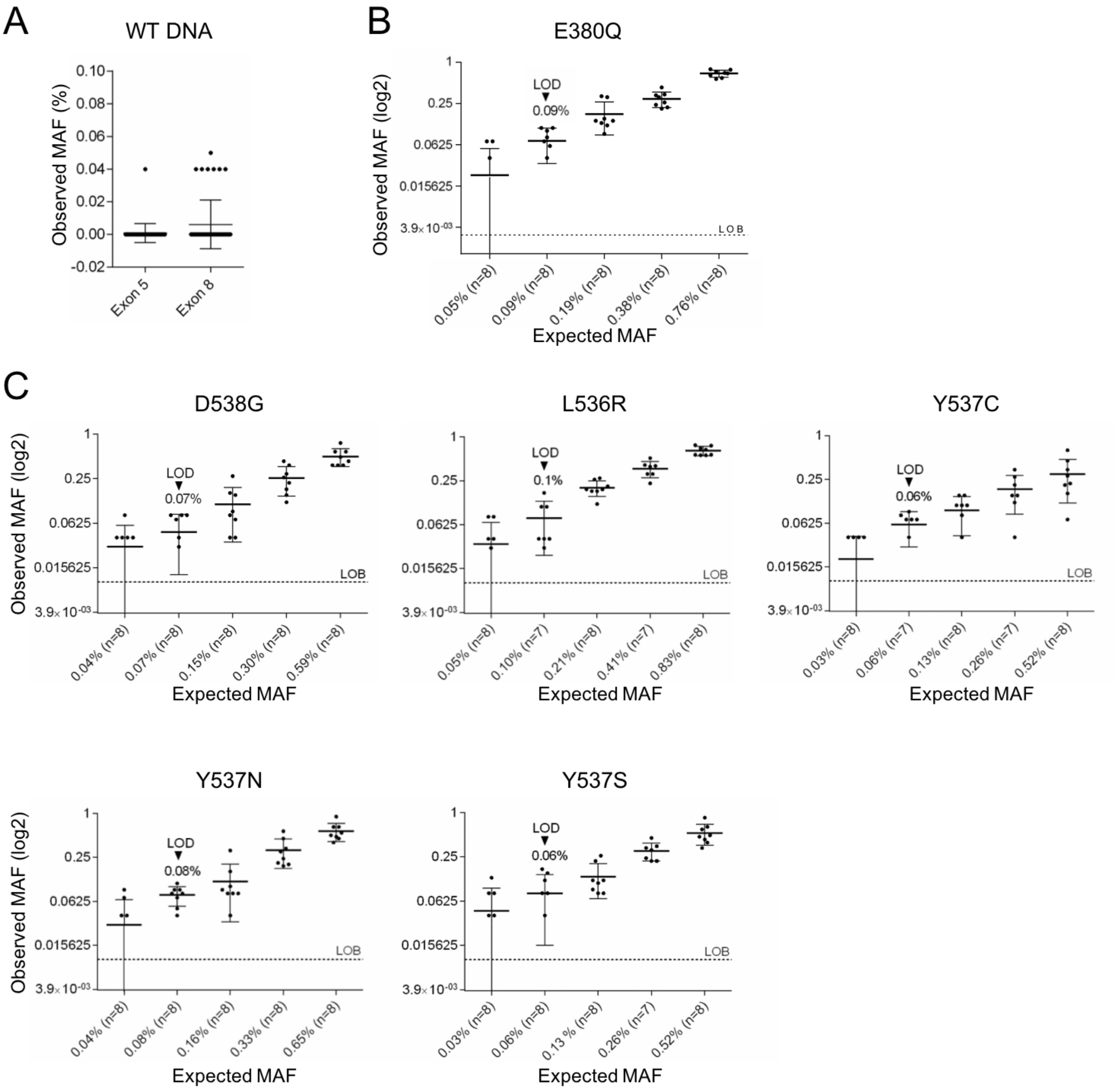


**Figure S1.** ***In vitro*** **performance of the simplex *ESR1*-ddPCR assays. A.** False positive events for exons 5 or 8 mutations observed from pure WT DNA tested with the E380Q assay or the Drop-off Ex8 assay. **B.** LOD estimation for exon 5 E380Q mutation tested with the E380Q assay. **C.** LOD estimation for exon 8 mutations D538G, L536R, Y537C, Y537N or Y537S tested with the Drop-off Ex8 assay. See method section for more details. LOB: limit of blank, LOD: limit of detection, estimated as the 95% CI of the mean false-positive calls.

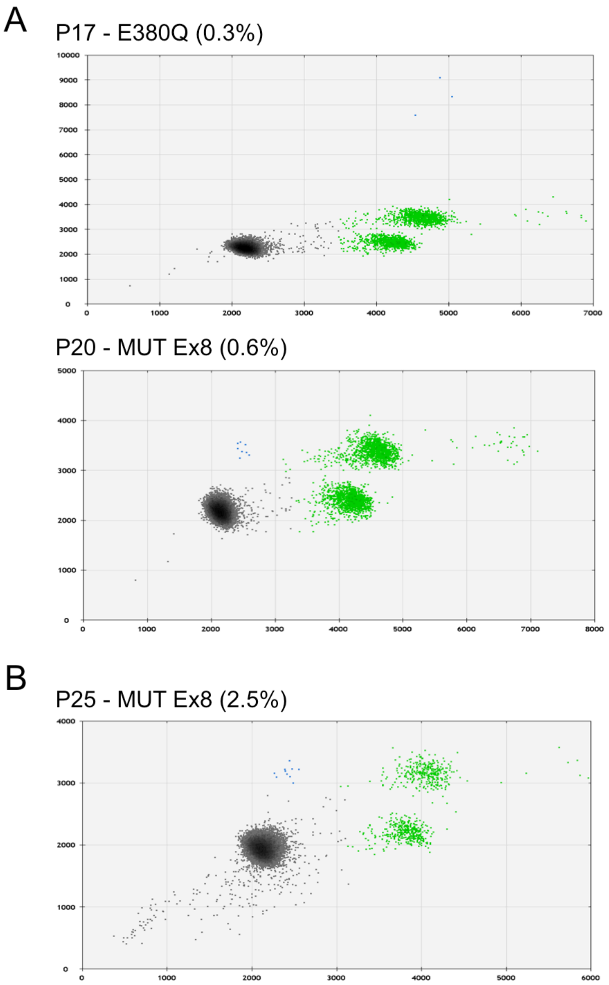


**Figure S2. Clinical samples with *ESR1* MAF <1% detected by ddPCR only. A.** ddPCR profiles for cases P-17 and P-20 not confirmed as mutant by NGS, as their MAFs were low (<1%). **B.** ddPCR profile for case P-25 not confirmed by NGS due to insufficient cfDNA for analysis. FAM and VIC intensities are shown in arbitrary units.

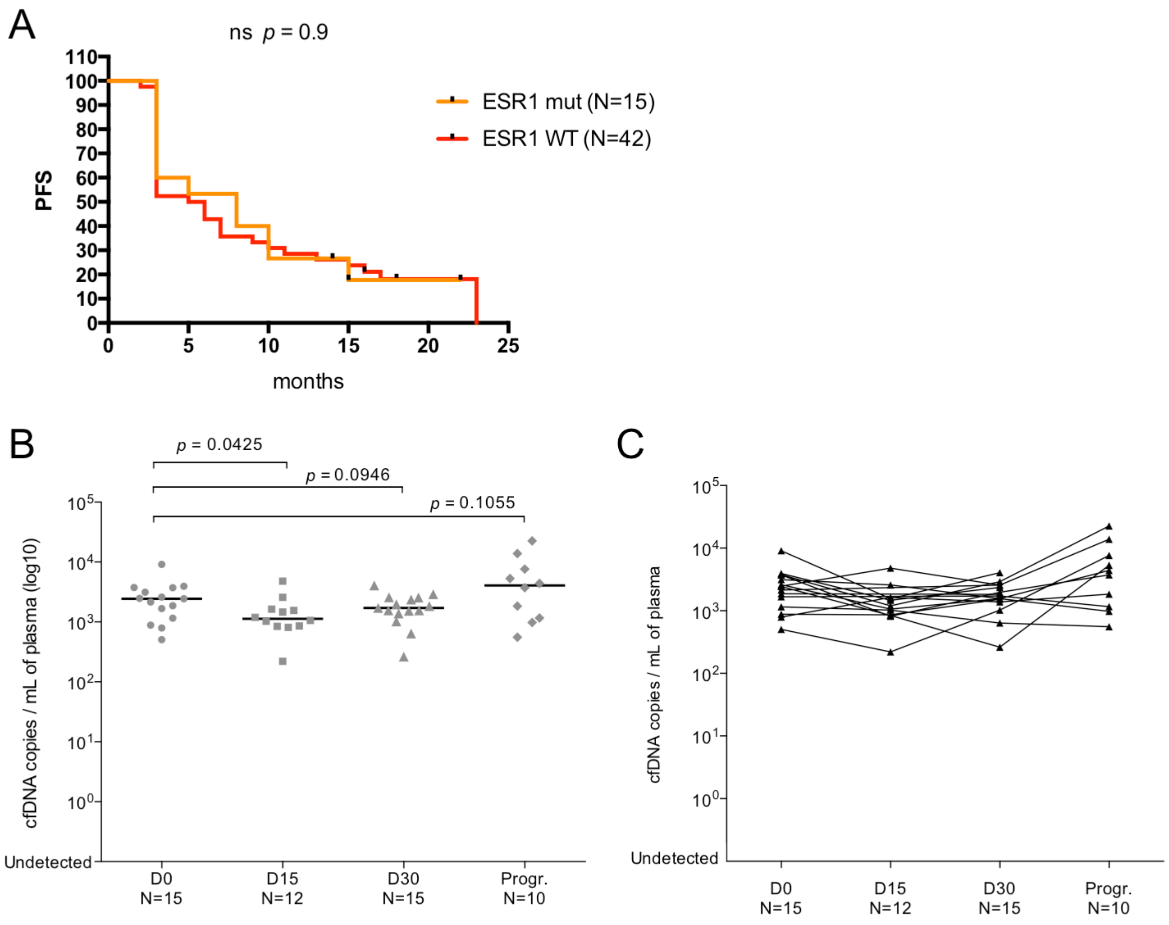


**Figure S3. *Circulating ESR1 mutant copies at baseline and its impact on PFS.* A.** Impact of *ESR1* status at baseline on PFS. **B.** Number of cfDNA copies detected per ml of plasma collected at the 4 time points during treatment follow-up (D0, N = 15; D15, N = 12; D30, N = 15; ToP, N = 10). **C.** cfDNA dynamics observed during treatment follow-up.

### Supplementary Tables

**Supplementary Table 1:** *ESR1* mutations targeted by the E380Q and drop-off Ex8 assays

**Supplementary Table 2:** Specificity tests for exon 5 and exon 8 assays in simplex and multiplex conditions.

**Supplementary Table 3:** Characteristics and ctDNA levels during treatment follow-up of patients carrying *ESR1* mutations
